# Genomics to aid species delimitation and effective conservation of the Sharpnose Guitarfish (*Glaucostegus granulatus*)

**DOI:** 10.1101/767186

**Authors:** Shaili Johri, Sam Fellows, Jitesh Solanki, Anissa Busch, Isabella Livingston, Maria Fernanda Mora, Anjani Tiwari, Asha Goodman, Adrian Cantu, Michael P. Doane, Megan Morris, Robert A. Edwards, Elizabeth A. Dinsdale

## Abstract

The Sharpnose Guitarfish (*Glaucostegus granulatus*) is one of fifteen critically endangered Rhino Rays which has been exploited as incidental catch, leading to severe population depletions and localized disappearances. Like many chondrichthyan species, there are no species-specific time-series data available for the Sharpnose Guitarfish that can be used to calculate population reduction, partly due to a lack of species-specific reporting as well as limitations in accurate taxonomic identification. We here present the first complete mitochondrial genome and partial nuclear genome of the species and the first detail phylogenetic assessment of the species. We expect that data presented in the current manuscript will aid in accurate species-specific landing and population assessments of the species in the future and will enable conservation efforts to protect and recover remaining populations.

## Introduction

The Sharpnose Guitarfish (*Glaucostegus granulatus*) is one of fifteen critically endangered Rhino Rays found in marine neritic and intertidal habitats of the northern Indian ocean, where it ranges from the Gulf of Oman and Persian Gulf to Myanmar [1]. This species, alongwith other Rhino Rays has been exploited as incidental catch, and this has led to severe population depletions and declines, and several localized disappearances [2]–[4]. Like many chondrichthyan species, there are no species-specific time-series data available for the Sharpnose Guitarfish that can be used to calculate population reduction [1]. This is due to a lack of species-specific reporting as well as limitations in accurate taxonomic identification. As a result current Red List assessments are based of off contemporary landings and catch rate datasets from range countries at varying levels of taxonomic resolution (eg. ‘Rhinobatids’ to ‘Guitarfishes’ to specific measurements for other sympatric species such as *Glaucostegus halavi* or *G. thouin* or *G. typus* or even Mobulid species). Based on these data, overall declines of >80% were estimated for Sharpnose Guitarfish populations throughout its range. However, it is known that, aggregated catches mask overfishing and local extinctions [5], underpinning the urgent need to enable species-specific reporting.

Second, Rhino Rays including the Sharpnose Guitarfish are proposed to be listed on Appendix II of the Convention for International Trade in Endangered Species (CITES), in August 2019 [6]. If the proposal is successful it will obligate nations to regulate all international exports of the species and require determination of sustainable and legal harvests of the species for exports to be permitted. Most fisheries that take the Sharpnose Guitarfish are intense and exploitative, poorly monitored, and unregulated for all practical purposes[1], [3]. Therefore, immediate protections through regulation of fishing and trade of the species are required to save the species from extinction. A crucial aspect of developing catch and trade regulations to protect remaining populations of *G.granulatus* and other Rhino Rays is to increase capacity for species specific reporting and accurate species identification. Such improvements are required to obtain the most accurate population estimates for each species from landing or catch data and to ensure absolute enforcement of trade laws in determining export quotas for each species under CITES.

Species identification is therefore vital to facilitate precise catch reporting and for enforcement of CITES regulations. However, identification of species alone is not sufficient to combat the complicated international trade networks involved in import-export of shark and ray products. International fin shipments change several hands and shipping containers during which they are also repackaged and relabeled, creating several loopholes and opportunities to mix illegally harvested fins (and other products) with legal harvests [7]. For example, traders may mix or replace legally harvested shark fins of species with those illegally harvested through ‘shark finning’ from a non-enforcing nation to abide by subsidy laws and launder illegal catch or to save on taxes by changing origin and destination labels. Indeed illegal global trading of endangered and CITES listed species has been reported by several studies [8]–[10]. For countries, that have laws against the import of fins from endangered species and against finning, it becomes critical to determine if a CITES listed species is being exported, and to determine if the fins are being sourced from a nation engaging in sustainable shark fisheries.

In order to determine source populations and country of origin, distinct populations of a commonly traded species need to be differentiated. In order to achieve the latter, genetic markers which allow differentiation of distinct geographical populations of a species are required. The current paradigm for control and regulation of trade relies solely on identification at the species level, in the best case scenario, with no consideration for distinct intraspecific populations. One of the main factors preventing species identification and incorporation of intraspecific differences in populations for trade regulations is the enormous data deficiency in genomic data for Chondrichthyes. Very few of the approximately 1200 Chondrichthyes have been assessed with respect to their population genetics and as a result it is impossible to differentiate distinct populations of a species. Overall, 46% of Chondrichthyes are data deficient, meaning there is insufficient information to determine their population and conservation status [11].

Molecular taxonomy of the Sharpnose Guitarfish and other Rhino Rays are limited and often based on a single genetic marker [12]. Whole mitochondrial genomes and nuclear genomic sequences can improve the resolution of chondrichthyan phylogenies [13], [14], and therefore provide more accurate species identification. In addition, whole mitochondrial genomes and nuclear markers provide important markers for population genetic analyses and identification of distinct populations, thus differentiating synonymous from structured populations [15]–[17].

We here report the first complete mitochondrial genome and partial nuclear genome of the Sharpnose Guitarfish, which was obtained by genome skimming of genomic DNA obtained from unprocessed fins slated for export. The mitogenome reported is the first complete sequence for *G.granulatus* and enabled us to improve species delimitation as well as phylogenetic relationships of taxa in the genus Glaucostegus. The mitochondrial and nuclear genomes could be used in future studies to identify SNPs and other molecular markers for assessment of population structure in the species. Last, acquiring population genetic data from a good sample size of *G. granulatus* in future studies, will enable quick determination of population estimates in different geographic areas, and these data can be used to prioritize target areas where in conservation efforts may be directed.

## Methods

DNA was extracted from a fin clip of an individual female guitarfish specimen (Figure 1) collected in Veraval, Gujarat, India and sequenced using the MinION sequencer (Oxford Nanopore Technologies) as outlined in Johri et al. 2019 [13]. Both Mitochondrial and nuclear genome fragments were obtained through Denovo assembly of the shotgun sequencing data from the sequencer in Geneious. The resulting mitochondiral genome sequence was annotated using the MitoAnnotator tool on the MitoFish website [18], and checked with ARWEN [19] and by comparison with annotated elasmobranch mitochondrial genomes from GenBank [20].

**Figure 1:**
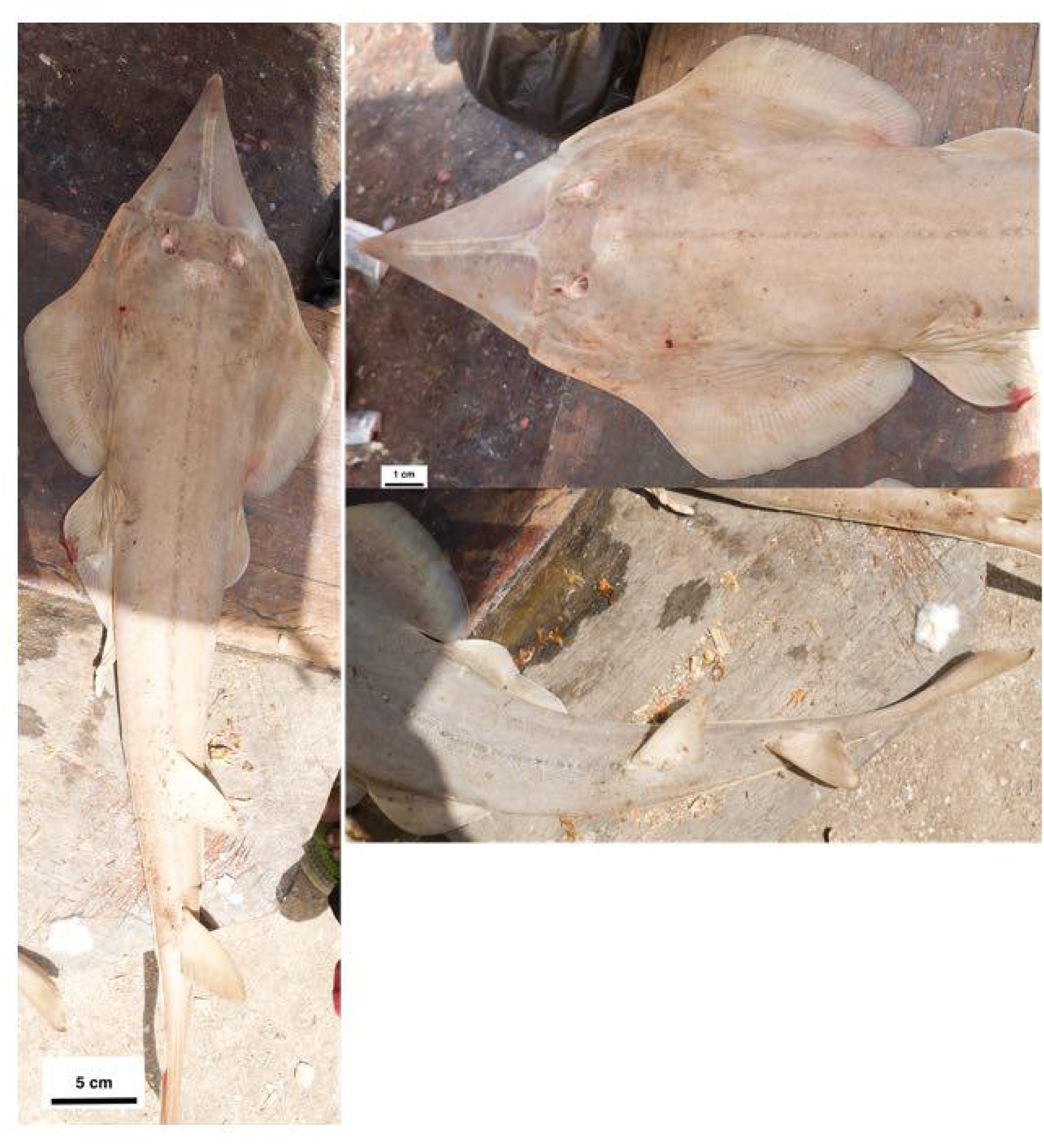
Dorsal view of specimen along the entire length (A), zoomed in views of the frontal (B) and caudal portions (C). Specimen identified as Glaucostegus granulatus using morphology.

Taxonomic identity of the sample was determined through morphological identification and phylogenetic assessment. Morphological identity was established through methods described in Rays of the World [21]. To assess phylogenetic placement of the specimen, gene trees were constructed using mitochondrial sequences for Cytochrome oxidase 1 (COX1), and NADH dehydrogenase subunit 2 (NADH2) obtained from GenBank [20] (Table S1). The genus Glaucostegus is not fully represented in any gene currently available on GenBank (five species each available for COX1, and NADH2). Genes were therefore analyzed separately and then in a concatenated matrix to assess congruence between phylogenies. Due to a large amount of missing data for the full COX1 gene, all COX1 sequences were trimmed to 717bp, the smallest available sequence. Multiple samples from *G.granulatus* and *G. thouin* were included to address monophyly of these two taxa since early analyses rendered them paraphyletic. All sequences were aligned using MUSCLE 3.8.31 [22] and PartitionFinder 2.1.1 [23] was used to identify optimal partitioning schemes and the best-fit model of molecular evolution, which was then used for all downstream analyses. In order to ensure robustness of the phylogenetic estimates, phylogenies were inferred in both Maximum Likelihood (IQ-Tree v1.6.10) [24] and Bayesian Inference frameworks, MrBayes v3.2.6 [25], [26]. For the IQ-Tree analyses, 100,000 ultrafast bootstrap replicates [27] were generated, beginning from 100 starting trees. MrBayes phylogenetic inference were run described in Johri et al. (2019) [13]. All analyses were run on XSEDE on CIPRES Scientific Gateway [28].

## Results and Discussion

The mitochondrial genome of *G.granulatus* (GenBank:) is 16,709 bp in length (Figure 2) and consists of 13 protein-coding genes (PCGs), 22 tRNA genes, 2 rRNA genes, and a non-coding control region (D-loop) (Table 1). The GC content is 40.0 % and the control region is 1014 bp long.

**Table 1:**

| **GENE NAME** | **TOPOLOGY** | **START-STOP** | **DIRECTION** |
| --- | --- | --- | --- |
| NADH dehydrogenase subunit 1 | CDS | 2876..3847 | + |
| NADH dehydrogenase subunit 2 | CDS | 4061..5106 | + |
| Cytochrome c oxidase subunit I | CDS | 5497..7053 | + |
| Cytochrome c oxidase subunit II | CDS | 7206..7896 | + |
| ATPase subunit 8 | CDS | 7972..8139 | + |
| ATPase subunit 6 | CDS | 8130..8813 | + |
| Cytochrome c oxidase subunit III | CDS | 8818..9603 | + |
| NADH dehydrogenase subunit 3 | CDS | 9675..10023 | + |
| NADH dehydrogenase subunit 4L | CDS | 10096..10392 | + |
| NADH dehydrogenase subunit 4 | CDS | 10386..11766 | + |
| NADH dehydrogenase subunit 5 | CDS | 11974..13815 | + |
| NADH dehydrogenase subunit 6 | CDS | complement(13811..14329) | - |
| Cytochrome b | CDS | 14403..15549 | + |
| 12S rRNA | rRNA | 72..1033 | + |
| 16S rRNA | rRNA | 1106..2799 | + |
| tRNA-Phe | tRNA | 1..71 | + |
| tRNA-Val | tRNA | 1034..1105 | + |
| tRNA-Leu | tRNA | 2800..2874 | + |
| tRNA-Ile | tRNA | 3849..3918 | + |
| tRNA-Gln | tRNA | complement(3918..3989) | - |
| tRNA-Met | tRNA | 3990..4060 | + |
| tRNA-Trp | tRNA | 5107..5176 | + |
| tRNA-Ala | tRNA | complement(5178..5246) | - |
| tRNA-Asn | tRNA | complement(5248..5321) | - |
| tRNA-Cys | tRNA | complement(5357..5424) | - |
| tRNA-Tyr | tRNA | complement(5427..5495) | - |
| tRNA-Ser | tRNA | complement(7057..7127) | - |
| tRNA-Asp | tRNA | 7128..7197 | + |
| tRNA-Lys | tRNA | 7897..7970 | + |
| tRNA-Gly | tRNA | 9605..9674 | + |
| tRNA-Arg | tRNA | 10024..10095 | + |
| tRNA-His | tRNA | 11767..11835 | + |
| tRNA-Ser | tRNA | 11836..11902 | + |
| tRNA-Leu | tRNA | 11902..11973 | + |
| tRNA-Glu | tRNA | complement(14331..14399) | - |
| tRNA-Thr | tRNA | 15550..15622 | + |
| tRNA-Pro | tRNA | complement(15625..15694) | - |
| control region | D-loop | 15695..16709 | + |

**Figure 2:**
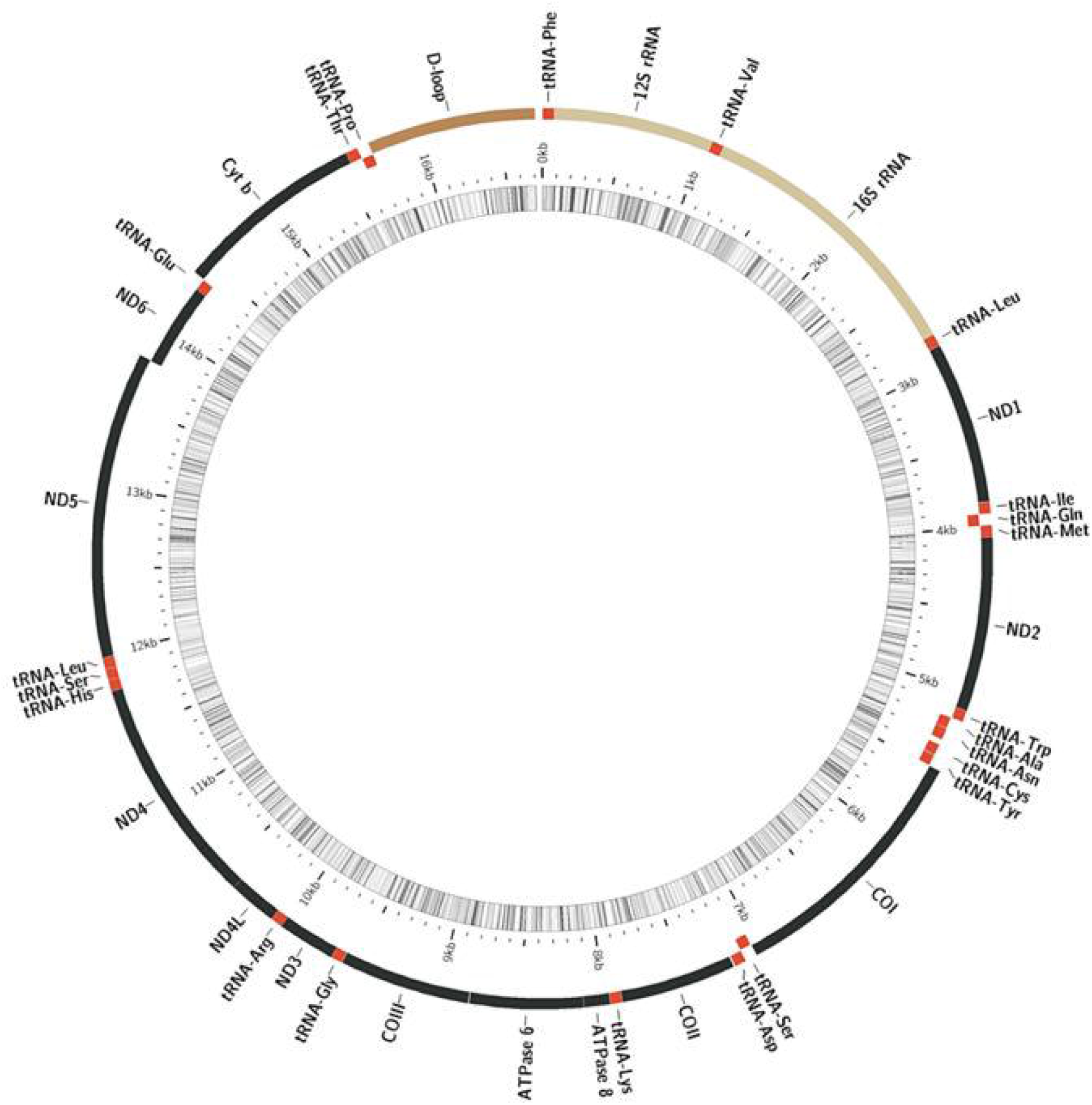
Annotated mitogenome sequence indicating gene position and topology of each loci, with CDS in black, tRNAs in red, rRNA in beige, and the control region in brown. Outer circle represents aligned genes with outermost representing the heavy strand and genes on the complementary l light strand depicted on the inside.

Phylogenetic analyses of Cytochrome oxidase 1 (COX1) sequences places the specimen within *Glaucostegus granulatus* with statistically significant support, however, much of the remainder of the tree has generally poor support in both the MrBayes and IQ-Tree analyses of COX1 (Figure S1A, S1B) due to poor data availability for wedgefishes. These analyses place G. thouin within *G. granulatus* with strong support, rendering *G. granulatus* paraphyletic at this marker. Our analyses of (ND2) eliminated the possibility of any resemblance of the specimen to G. halavi, which was not present in the COX1 dataset due to sequence unavailability. ND2 runs also recovered poor support throughout the remaining phylogenetic tree (Figure S2A, S2B) using both MrBayes and IQ-Tree analyses.

Phylogenetic analyses of the concatenated matrix again placed the specimen within *G. granulatus*, and rendered *G. granulatus* paraphyletic with strong support. While MrBayes recovered high support throughout the in-group taxa (Figure 3A), IQ-Tree analysis recovered moderate bootstrap support throughout the remainder of the tree (Figure 3B). Furthermore, our MrBayes analysis placed one *G. thouin* sample as sister to *G. typus* rather than in the *G.granulatus* + thouin clade (Figure 3A).

**Figure 3A:**
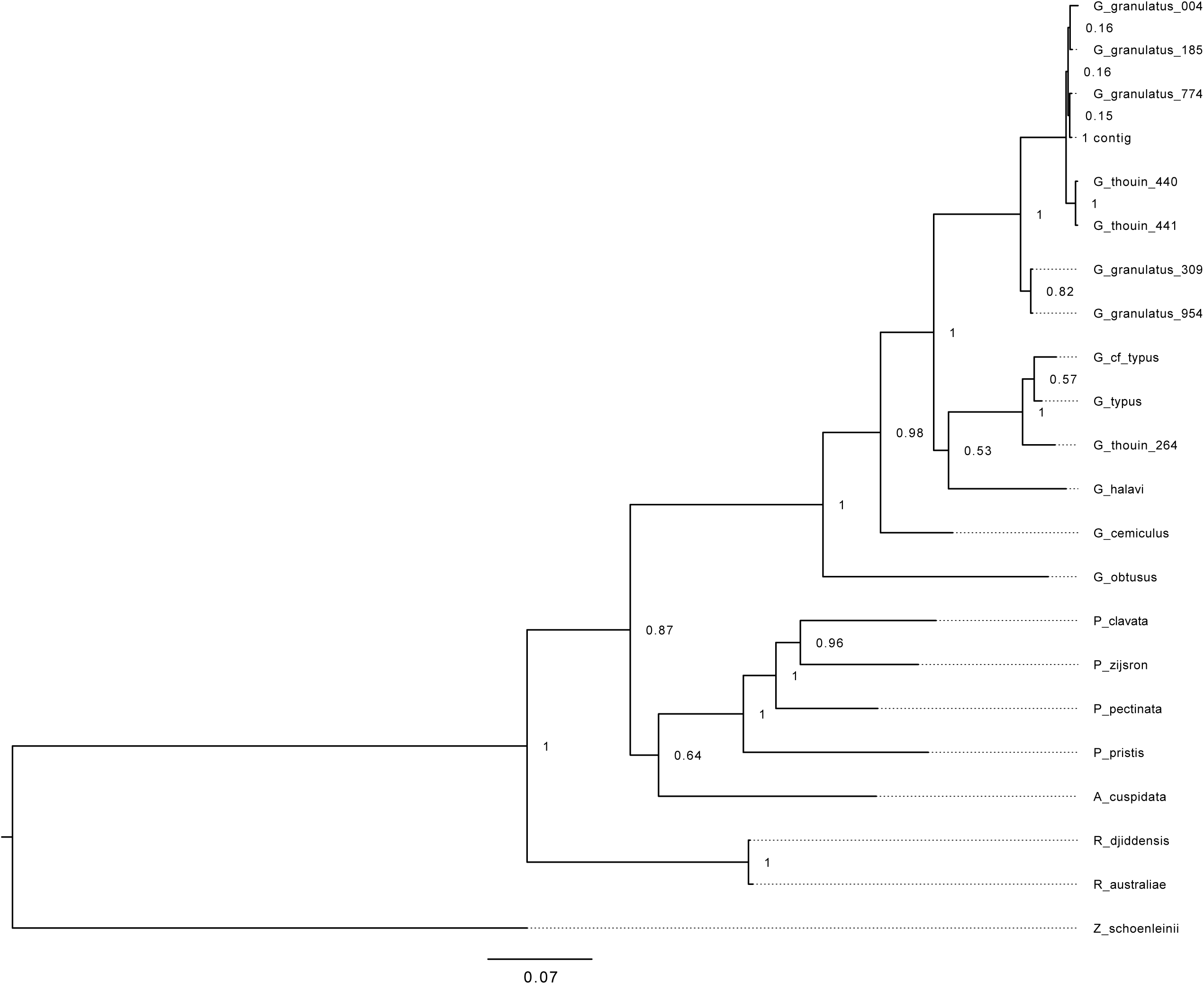
Bayesian estimate of relationships among representative taxa from orders Rhynchobatiformes, and Pristiformes. Bayesian phylogenetic estimates determined from concatenated sequences of COI and NADH2 mitochondrial genes from 21 species with *Zanobatos schoenleini* as outgroup. The unknown sample clusters with *G.granulatus*. Numbers at nodes are posterior probabilities.

**Figure 3B:**
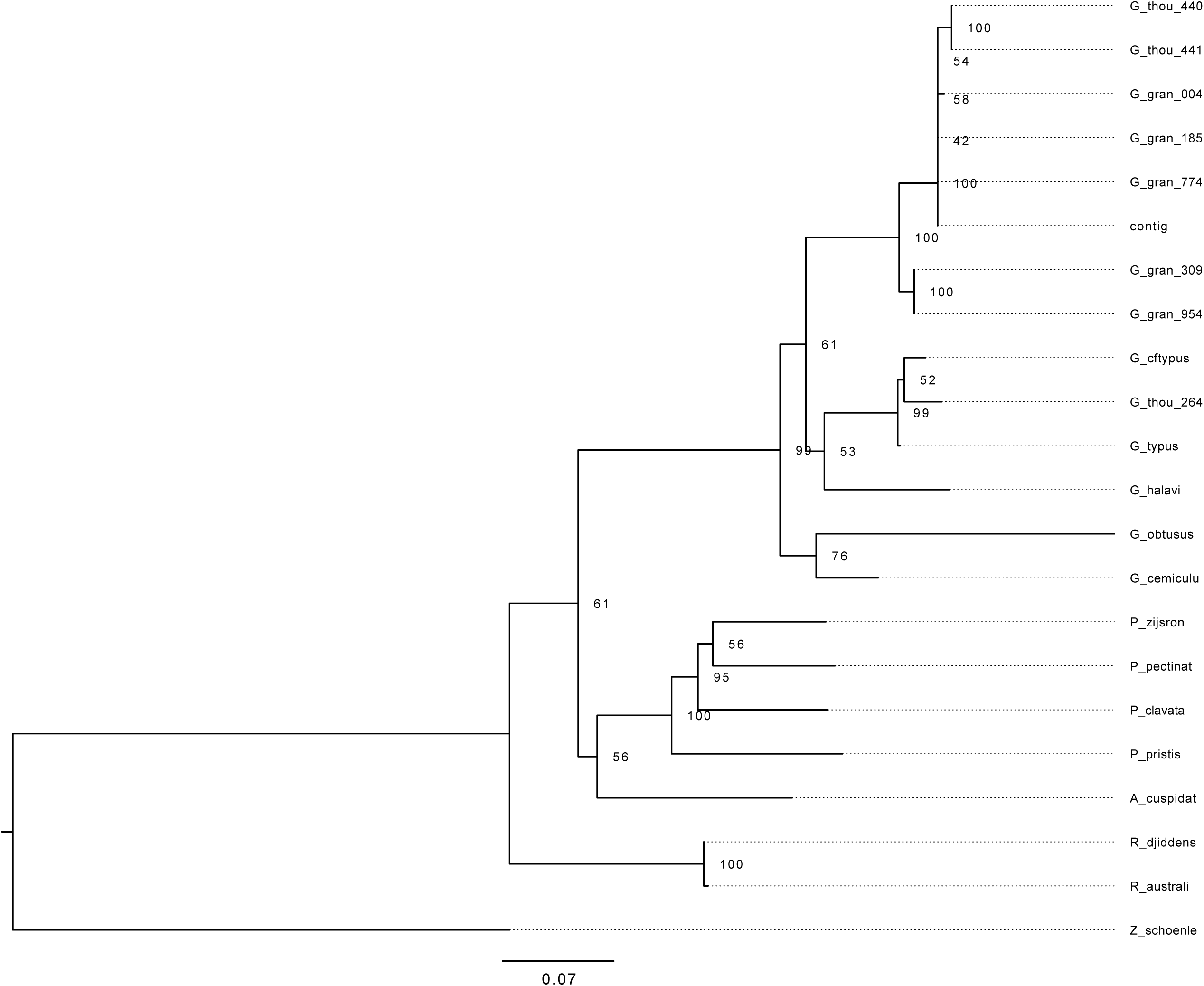
Maximum-likelihood estimate of relationships among Rhyncobatiformes and Pristiformes using concatenated mitochondrial protein coding sequences. Maximum-likelihood phylogenetic estimates determined from concatenated sequences of COI and NADH2 mitochondrial genes from 21 species with *Zanobatos schoenleini* as outgroup. The unknown sample clusters with *G.granulatus*. Numbers at nodes are bootstrap support values.

The current report presents the first complete mitochondrial genome and partial nuclear genome of *G.granulatus*. These genomic data will significantly aid identification and assessment of population and conservation status of the species. Overall, the genomic sequences and the analyses presented in this report are a step forward in reducing the data deficiency of the species and in general for the genus Glaucostegus.

We present the most extensively-sampled published phylogeny of the genus Glaucostegus by including six species, two of which were represented by multiple samples. Our phylogenetic analyses confidently place the specimen under consideration within *G. granulatus*, matching its taxonomic classification to *G. granulatus* using morphological parameters. While our analyses support the monophyly of Glaucostegus and broadly find similar relationships as previous work [29], [30], resolution of the full phylogenetic tree is limited by lack of available data. Full resolution of relationships among the Rhinobatid species will require additional genetic or genomic datasets and inclusion of taxa which currently lack sequence data (*G. microphthalamus and G. spinosus*).

In addition to phylogenetic placement of our specimen, our analyses places two previously reported samples of *G. thouin* within *G. granulatus* and one other as sister to *G. typus* as also reported by Aschliman, N. 2011 [29], suggesting undescribed diversity or complex demographic processes within Glaucostegus. Further sampling with additional markers will be required to resolve these relationships.

## Supporting information

Supplementary Information_Mitogenome

Supplementary Information_PartialGenome

