## Supplementary figures and images for "Genomics to aid species delimitation and effective conservation of the Sharpnose Guitarfish (*Glaucostegus granulatus*)"

### Supplementary Figure S1A

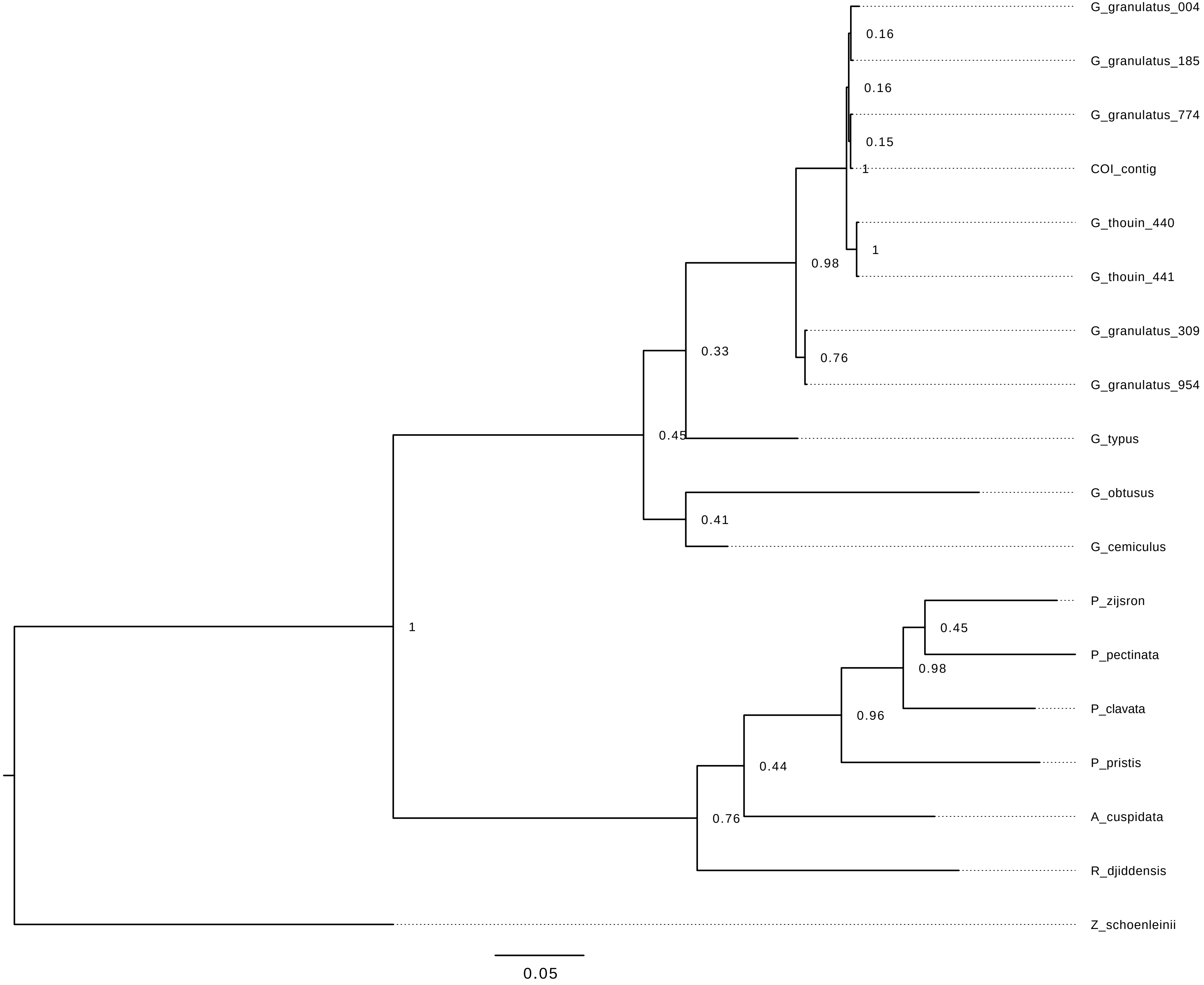

### Supplementary Figure S1B

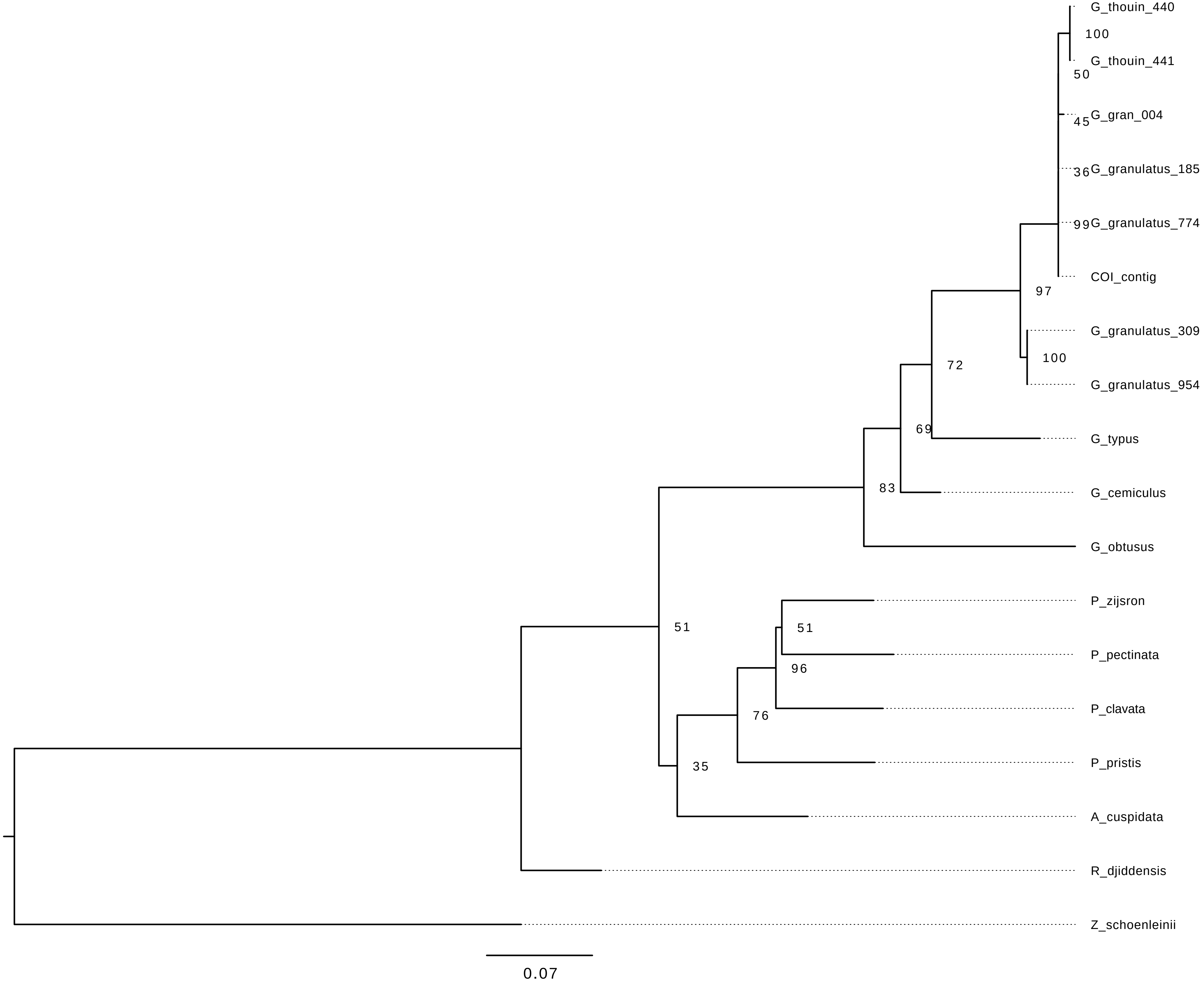

### Supplementary Figure S2A

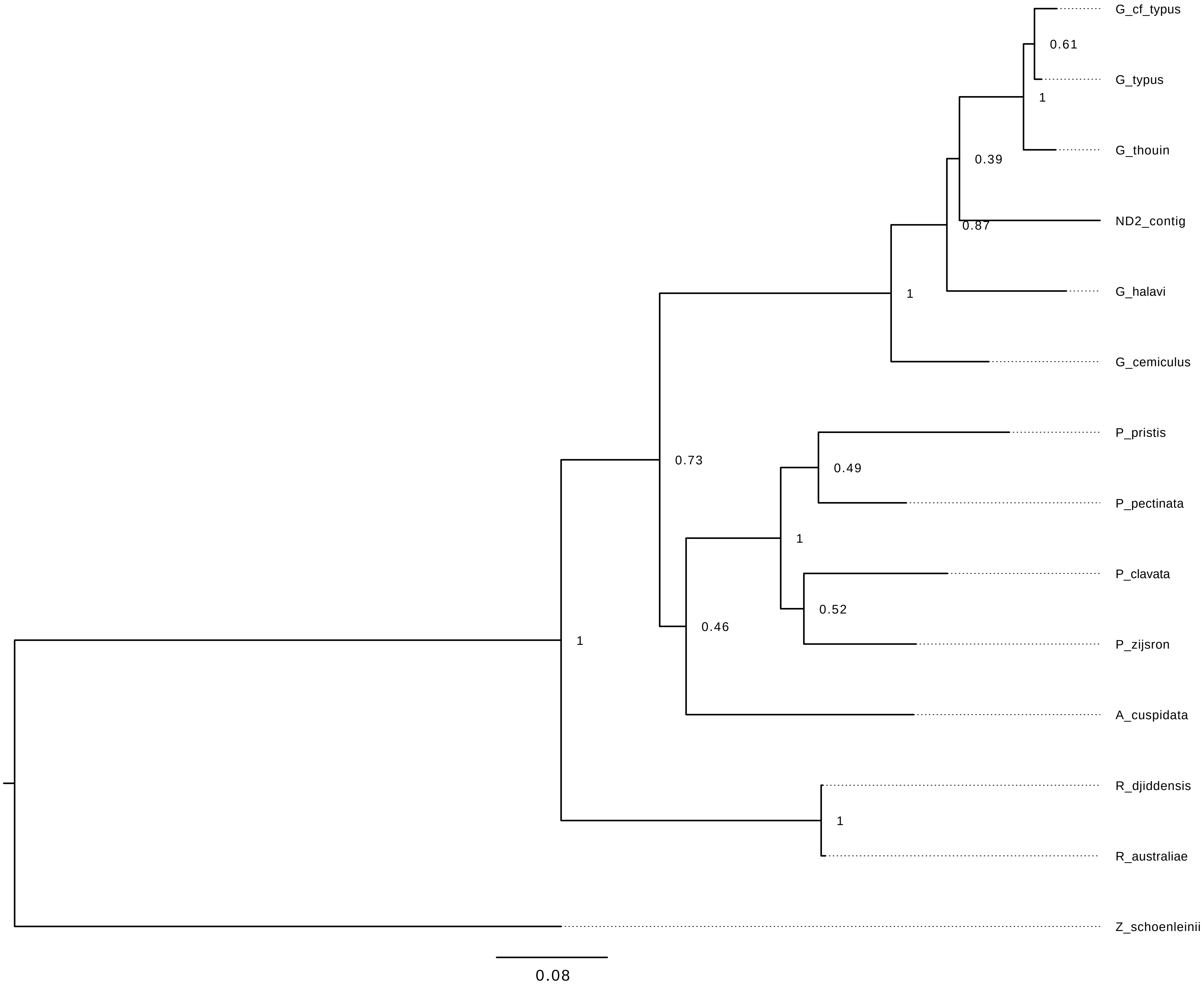

### Supplementary Figure S2B

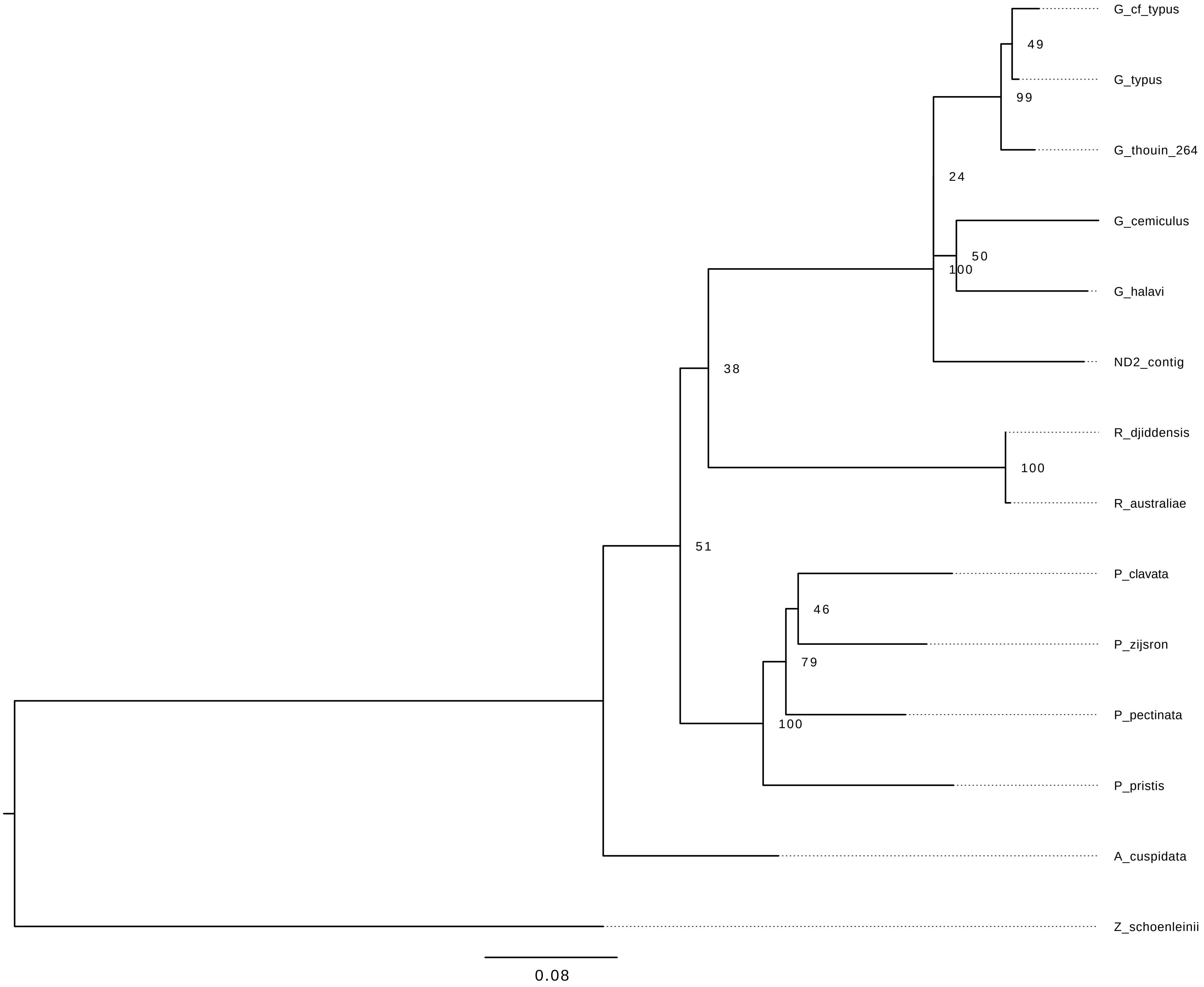
